# SARS-CoV-2 inactivation by human defensin HNP1 and retrocyclin RC-101

**DOI:** 10.1101/2021.05.27.445985

**Authors:** Elena Kudryashova, Ashley Zani, Geraldine Vilmen, Amit Sharma, Wuyuan Lu, Jacob S. Yount, Dmitri S. Kudryashov

## Abstract

Severe acute respiratory syndrome coronavirus (SARS-CoV)-2 is an enveloped virus responsible for the COVID-19 respiratory disease pandemic. While induction of adaptive antiviral immunity via vaccination holds promise for combatting the pandemic, the emergence of new potentially more transmissible and vaccine-resistant variants of SARS-CoV-2 is an ever-present threat. Thus, it remains essential to better understand innate immune mechanisms that are active against the virus. One component of the innate immune system with broad anti-pathogen, including antiviral, activity is a group of cationic immune peptides termed defensins. The defensins’ ability to neutralize enveloped and non-enveloped viruses and to inactivate numerous bacterial toxins correlate with their ability to promote the unfolding of thermodynamically pliable proteins. Accordingly, we found that human neutrophil a-defensin HNP1 and retrocyclin RC-101 destabilize SARS-CoV-2 Spike protein and interfere with Spike-mediated membrane fusion and SARS-CoV-2 infection in cell culture. We show that HNP1 binds to Spike with submicromolar affinity. Although binding of HNP1 to serum albumin is more than 20-fold weaker, serum reduces the anti-SARS-CoV-2 activity of HNP1. At high concentrations of HNP1, its ability to inactivate the virus was preserved even in the presence of serum. These results suggest that specific a- and 8-defensins may be valuable tools in developing SARS-CoV-2 infection prevention strategies.

## INTRODUCTION

COVID-19 imposes an extraordinary health threat whose scale and severity can be compared only to that of the Spanish flu pandemic, which raged over a hundred years ago. The COVID-19 pandemic vividly exposed the weakness of the globalized world to new zoonotically transmitted viruses, which emerge with alarming regularity (e.g., 2003 Severe Acute Respiratory Syndrome coronavirus SARS-CoV ^1^, 2009 influenza A H1N1 ^2^, 2012 Middle East Respiratory Syndrome coronavirus MERS-CoV ^3^). Although several effective SARS-CoV-2 vaccines have been generated in an unprecedently short time frame, high levels of transmissibility and mortality of this virus [reviewed in ^4^], hesitancy towards vaccination from a large fraction of the world population, and the emergence of vaccine-resistant variants require additional development of non-vaccine therapies against SARS-CoV-2.

Interestingly, despite the high mortality rate, as much as 42% of infected individuals are asymptomatic carriers of SARS-CoV-2 ^5-7^ suggesting that the virus can be effectively controlled by the human innate immune system ^8^. Recently, pro- and antiviral activities of interferon-induced transmembrane proteins (IFITMs) in SARS-CoV-2 infection have been described ^9-11^. Likewise, other interferon-stimulated genes, such as CH25H, Ly6E, Tetherin, and ZAP have been demonstrated to have inhibitory effects on SARS-CoV-2 replication in vitro ^12-16^. Another important and promising class of innate immunity effectors is defensins. These peptides are active against several viruses, including human immunodeficiency virus (HIV), herpes simplex virus (HSV), influenza, and SARS-CoV ^17, 18^.

Based on their structural characteristics, mammalian defensins are separated into three subfamilies called a-, -, and 8-defensins ^19-22^. Humans express a- and -defensins, while cyclic 8-defensins are produced only by non-human primates ^23^. Although human 8-defensin genes are transcribed, a premature stop codon precludes their translation ^24^. Intriguingly, human cells retain the ability to produce the cyclic 8-defensin peptides upon transfection of the synthetic “humanized” 8-defensin genes called retrocyclins (RC) ^25^. Potent antiviral and anti-bacterial activity ^23, 26-28^ in combination with low cytotoxicity and exceptional stability ^29, 30^ identified RCs as promising topical microbicides ^30-32^.

α-defensin human neutrophil peptides 1-4 (HNP1-4) are highly homologous dimeric cationic amphiphilic peptides ^33^, among which HNP1 is the most abundant. HNP1 shows potent anti-bacterial ^34-37^, anti-fungal ^38, 39^, and antiviral activities against enveloped and non-enveloped viruses (reviewed in ^40^). Antiviral mechanisms of defensins (reviewed in ^41^) are multifaceted, including direct targeting of exposed proteins on viral envelopes and capsids, disruption of viral fusion, inhibition of post-entry processes, or affecting receptors on the host cell surface. Such remarkable versatility is partially explained by the ability of defensins to disrupt membranes ^34^, but also by their ability to induce unfolding of marginally stable proteins, particularly bacterial toxins and viral proteins ^42-44^. Susceptibility of target proteins to defensin-induced destabilization, unfolding, and ultimately inactivation is reliant on their conformational plasticity, which often correlates with thermolability (reviewed in ^45^). As evolutionary derivatives of a-defensins ^23^, 8-defensins (and retrocyclins, in particular) mainly recapitulate their activity ^46^. Defensins are also recognized as chemokines that modulate the immune response by binding to cell surface receptors and interfering with cellular signaling ^41^. According to a recent study, enteric human a-defensin HD5 binds to human angiotensin-converting enzyme 2 (a.k.a. hACE2 receptor) on enterocytes obstructing SARS-CoV-2 Spike binding site, which results in inhibition of SARS-CoV-2 Spike pseudovirion infection ^47^. Encouraged by this report, we examined the effects of a neutrophil a-defensin, HNP1, and retrocyclin, RC-101, on SARS-CoV-2 infection.

## RESULTS AND DISCUSSION

### HNP1 and RC-101 inhibit SARS-CoV-2 Spike-mediated membrane fusion and pseudotyped virus infection

SARS-CoV-2 homotrimeric Spike glycoprotein binds to the hACE2 receptor on the host cell, mediating membrane fusion and virus entry ^48^. Being exposed on the surface of virions, Spike is the main target for neutralization of the virus by antibodies. Membrane fusion activity of Spike enables fusion of the mammalian cells ectopically producing Spike with hACE2-positive cells into multicellular syncytia ^49^. Such syncytia represent a convenient model to study Spike/hACE2 interaction and to test different agents in disrupting this interaction. Another approach that does not require stringent biosafety level 3 (BSL3) conditions is to use replication-restricted pseudotyped viruses encoding SARS-CoV-2 Spike.

We explored both options to test whether HNP1 and RC-101 can interfere with the SARS-CoV-2 Spike interaction with host cells. Co-transfection of human osteosarcoma U2OS cells with Spike and mCherry promoted their fusion with ACE2-producing human lung Calu-3 cells, leading to multinuclear mCherry-positive syncytia (Fig. 1A). In the absence of serum, the addition of HNP1 or RC-101 to the co-culture of Spike- and ACE-2-positive cells strongly inhibited syncytia formation (Fig. 1A,B). Under these conditions, both defensins induced the formation of Hoechst-negative precipitate visible only in the phase-contrast channel, which excludes microbial contamination. In agreement with previous reports ^37, 50^, serum abrogated the defensin’s effect (Fig. 1C).

**Figure 1.**
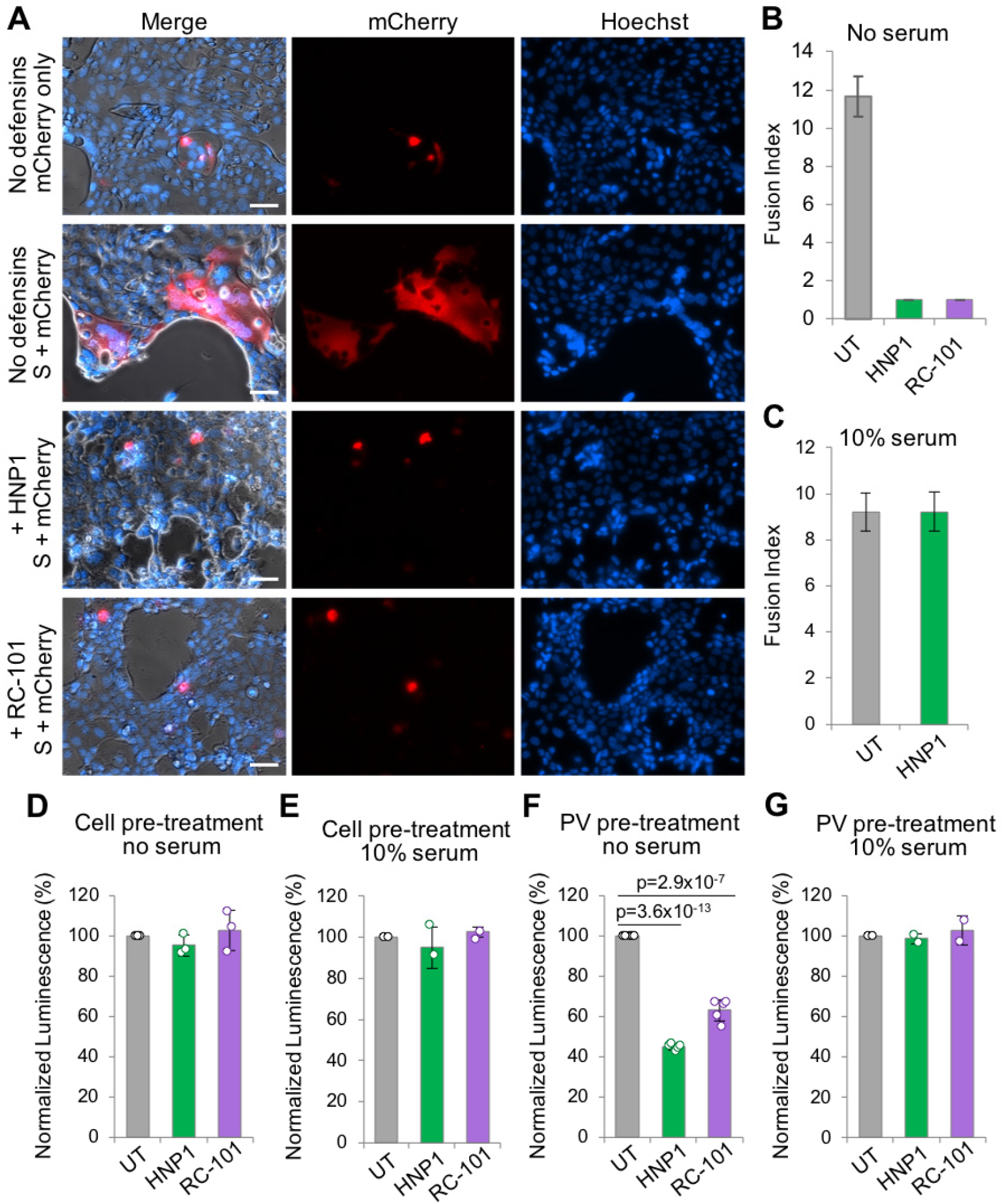
HNP1 and RC101 inhibit SARS-CoV-2 Spike-pseudotyped viral infection. **(A-C)** U2OS cells co-transfected with SARS*-*CoV*-*2 Spike (S) and mCherry were mixed with Calu*-*3 cells, co*-*cultured for 24 h in the absence or presence of 40 µM HNP1 or RC-101, and imaged by fluorescent microscopy to visualize mCherry and nuclei stained with Hoechst dye. **(A)** Representative images of the cells co-cultured in the absence of serum; scale bars are 50 µm. mCherry*-*positive cell syncytia were formed upon Spike expression in the absence of serum and defensins. **(B, C)** Fusion indices were quantified as an average number of nuclei per individual mCherry*-*positive syncytium (66-77 individual syncytia were quantified) in the absence **(B)** or presence of 10% serum **(C)**. Data is presented as mean ± SE. UT - untreated cells without defensins. **(D-G)** Infection efficiency of SARS-CoV-2 Spike-pseudotyped viruses encoding Nluc luciferase was monitored by luminescence assays. **(D, E)** H1299 cells were either untreated (UT) or pretreated with 50 µM HNP1 or RC-101 before the addition of the pseudoviruses in the absence **(D)** or presence of 10% serum **(E). (F, G)** H1299 cells were transduced by pseudoviruses, which were either untreated (UT) or pretreated with 50 µM HNP1 or RC101 before the addition to the cells in the absence **(F)** or presence of 10% serum **(G)**.

Next, human lung H1299 cells were infected with HIV-based reporter virus-like particles pseudotyped with SARS-CoV-2 Spike. Pseudovirus infection was monitored using a secreted NanoLuc luciferase (Nluc) reporter. Pre-treatment of the cells with HNP1 or RC-101 prior to the infection did not affect the pseudovirus infection efficiency (Fig. 1D,E). In contrast, pre-incubation of the pseudovirus particles with the defensins for 1 hour significantly inhibited the infection (Fig. 1F). These results suggest that HNP1 and RC-101 can interfere with the viral infection by acting upon the viral particle rather than influencing the host’s molecules (e.g., hACE2 receptor), as it has been suggested for enteric defensin HD5 ^47^. As with the Spike-mediated fusion experiments, serum had a strong negative influence on the inhibitory ability of HNP1 and RC-101 (Fig. 1G).

### HNP1 binds to and destabilizes SARS-CoV-2 Spike protein

The results from the SARS-CoV-2 Spike-mediated membrane fusion and pseudovirus infections prompted us to test whether HNP1 can directly interact with Spike protein. Fluorescence anisotropy revealed that Cy5-HNP1 strongly binds to a recombinant SARS-CoV-2 Spike with submicromolar affinity (K_d_ = 146 ± 18 nM; Fig. 2A). Given the inhibitory effect of serum, we checked whether Cy5-HNP1 binds serum albumin, the most abundant and highly interactive serum protein. Indeed, Cy5-HNP1 was able to bind to bovine serum albumin (BSA), albeit with over 20-fold lower affinity (K_d_ = 3.4 ± 0.3 µM; Fig. 2B), pointing to its much higher selectivity towards Spike.

**Figure 2.**
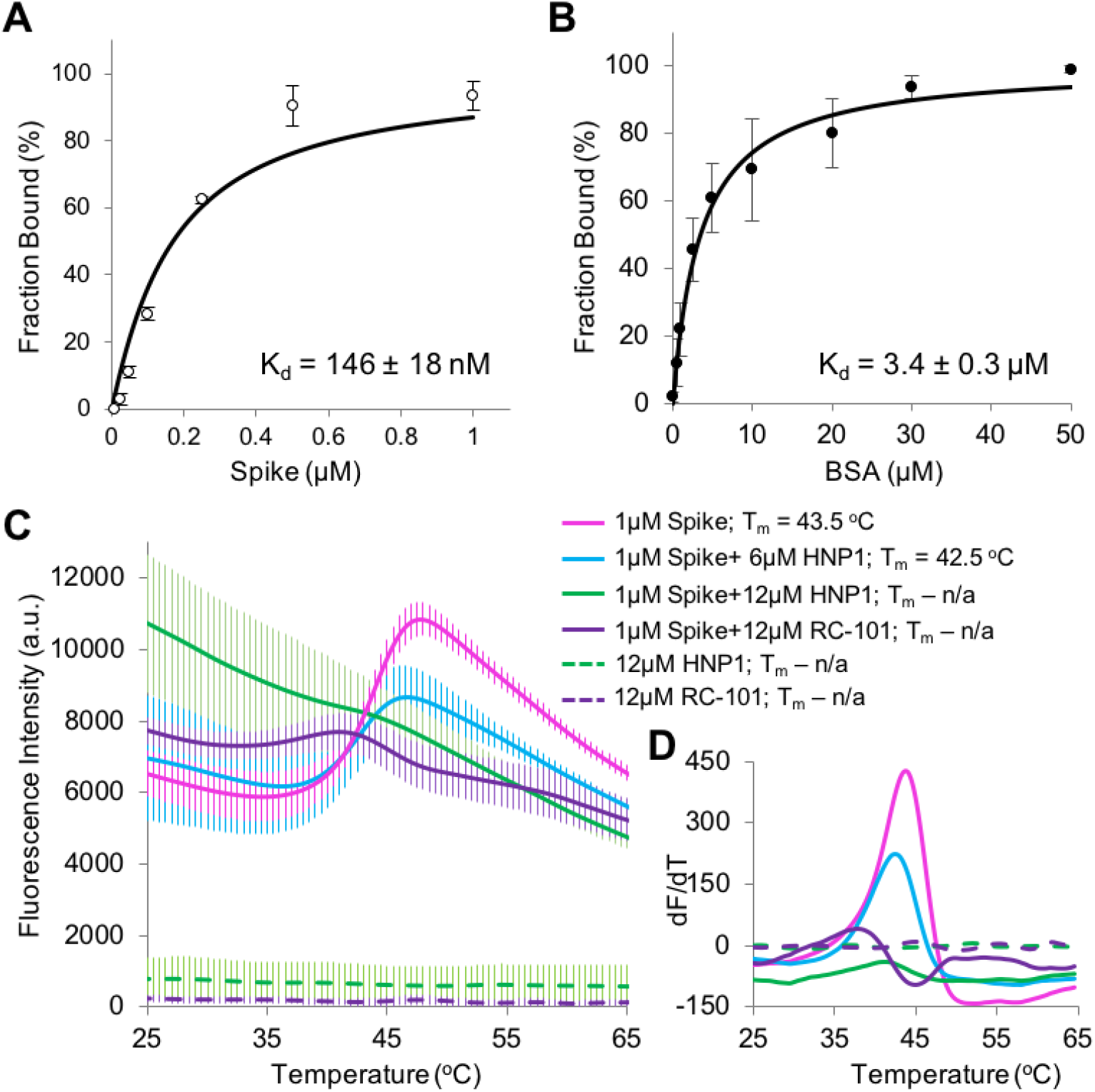
HNP1 and RC-101 destabilize SARS-CoV-2 Spike protein. **(A, B)** Binding of 50 nM Cy5-HNP1 to SARS-CoV-2 Spike **(A)** or BSA **(B)** was monitored by fluorescence anisotropy. Data from two independent experiments performed in duplicates is presented as mean ± SE. The K_d_s were calculated by fitting the data to the binding isotherm equation (see Methods). **(C, D)** Stability of SARS-CoV-2 Spike protein in the absence and presence of HNP1 and RC-101 was assessed by DSF.**(C)**Data from three independent experiments is presented as mean ± SE. **(D)** First derivatives of the fluorescence versus time traces are shown. Melting temperatures (T_m_) were calculated as the infliction points of the derivative curves; n/a - not applicable.

Defensins inactivate various structurally unrelated pathogen effector proteins such as bacterial toxins and viral proteins ^42, 43, 46^. A common trait that unites these targets is their marginal thermodynamic stability dictated by the need for transitioning through dramatic conformational perturbations required for passing through narrow pores (i.e., for bacterial toxins) or fusing viral and host membranes (i.e., viral fusion proteins) ^44, 45^ For viral proteins, conformational plasticity is also dictated by a higher ability of such “loosely” folded proteins to tolerate high mutational loads. SARS-CoV-2 Spike is subject to both types of evolutionary pressure and, as such, is anticipated to be prone to inactivation by defensins. Therefore, we assessed the effects of HNP1 and RC-101 on the stability of Spike using differential scanning fluorimetry (DSF) ^51^. Spike protein was indeed destabilized at micromolar concentrations of HNP1 and RC-101 (Fig. 2C,D) as implied from the impaired melting profile with a higher signal at low temperatures and a loss of the peak at its normal position. Such behavior is characteristic of chemical denaturation as it reports the fluorophore binding to hydrophobic residues of Spike exposed at temperatures below the melting point of thermal denaturation.

### HNP1 inhibits SARS-CoV-2 infection with low-micromolar efficiency

We next evaluated whether the inhibitory effects of HNP1 and RC-101 observed in the pseudovirus infections are also observed with the genuine SARS-CoV-2 virus in a BSL3 controlled environment. Vero E6 cells susceptible to SARS-CoV-2 infection ^52^ were infected with SARS-CoV-2, and the efficiency of infection was measured by flow cytometry staining for the viral nucleocapsid (N) protein. In the first set of experiments (Fig. 3), the virus was pre-incubated with the defensins (50 µM final concentration) in the absence of serum. Under these conditions, SARS-CoV-2 infectivity was blocked by HNP1 and strongly inhibited by RC-101 (Fig. 3A,B). Serum significantly reduced the ability of HNP1 to inhibit the infection in a concentration-dependent manner (Fig. 3C,D).

**Figure 3.**
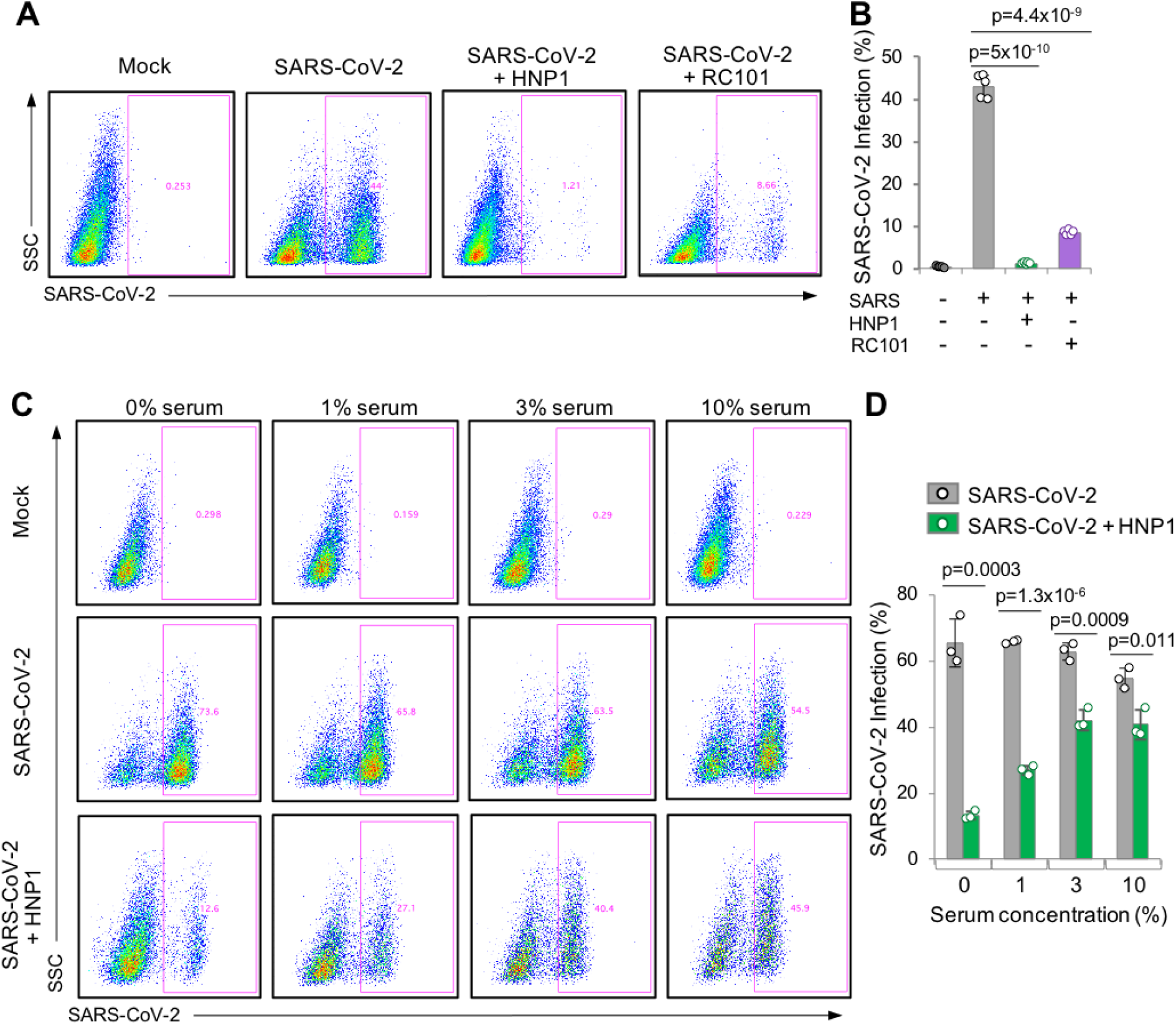
Pre-incubation of SARS-CoV-2 viruses with HNP1 or RC101 inhibits infection in a serum-dependent manner. Vero E6 cells were infected with SARS*-*CoV*-*2 and analyzed for infection by flow cytometry. **(A, B)** Cells were infected with the SARS-CoV-2 viruses pretreated with 50 µM HNP1 or RC101 in the absence of serum. **(C, D)** Cells were infected with the SARS-CoV-2 viruses pretreated with 25 µM HNP1 with increasing serum concentrations. **(A, C)** Representative example plots are shown for each condition. **(B, D)** Graphs depict mean infection percentage measurements from experiments conducted in 5 or 3 replications. Error bars represent the SD; p values determined by Student’s t-test are indicated.

Next, we found that pre-incubation is not essential for the virus neutralization, as HNP1 potently inhibited SARS-CoV-2 infection when added to the cells immediately after the virus (Fig. 4A,B). Without serum, the inhibitory concentration (IC_50_) of HNP1 was 10.3 ± 0.9 µM (Fig. 4C-E). Notably, HNP1 strongly inhibited the SARS-CoV-2 infection even in the presence of 10% serum, providing that the peptide was present at sufficiently high concentration (e.g., 43 µM; Fig. 4B).

**Figure 4.**
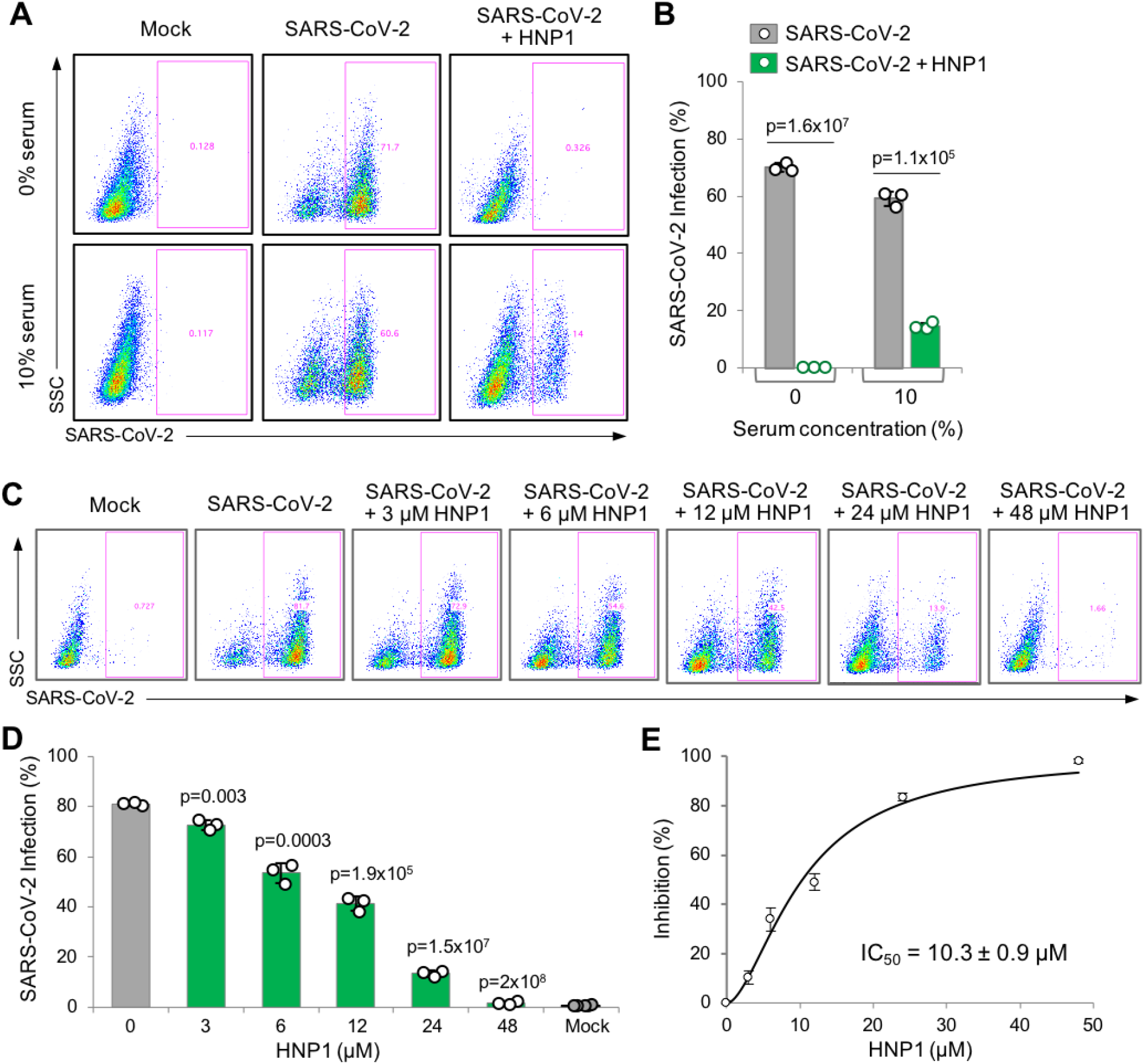
HNP1 inhibits SARS-CoV-2 infection with low-micromolar efficiency without pre-incubation. Vero E6 cells were infected with SARS*-*CoV*-*2 and analyzed for infection by flow cytometry. **(A, B)** Cells were infected with SARS-CoV-2 in the presence of 43 µM HNP1 in the absence and presence of 10% serum. **(C-E)** Cells were infected with SARS-CoV-2 in the absence of serum with increasing concentrations of HNP1 to calculate IC_50_ **(E)** by fitting the data from **(D)** to Hill’s equation (see Methods). **(A, C)** Representative example plots are shown for each condition. **(B, D)** Graphs depict mean infection percentage measurements from experiments conducted in three replications. Error bars represent the SD; p values determined by Student’s t-test are indicated.

To summarize, high concentrations of a-defensin HNP1 and 8-defensin (retrocyclin) RC101 are capable of neutralizing SARS-CoV-2 infection. Although even higher levels of HNP1-4 are likely achievable near the neutrophils immediately upon release of the defensins, they are quickly diluted, and their concentration in body fluids is much lower. For example, the level of HNP1-4 in the nasal aspirates of children affected by adenovirus was about 220 ng/mL (∼0.07 µM) ^53^. Even though the inhibitory effects of the HNP1 and RC101 against SARS-CoV-2 are diminished by serum, the defensins should be explored as topical (nasal) antiviral agents. In particular, retrocyclins have been proposed as topical microbicides to prevent sexually transmitted infections caused by HIV-1 ^54, 55^. Notably, owing to their cyclic nature, high stability, and resistance to proteolysis, retrocyclins are particularly promising. RCs have low toxicity in cell cultures and in vivo, are well tolerated in animal models, and are non-immunogenic in chimpanzees ^56, 57^. Together our data suggest that HNP1 and RC-101 should be further evaluated as candidates for developing topical anti-COVID and broad-range antiviral drugs.

## MATERIALS AND METHODS

### Peptides, proteins, and DNA constructs

HNP1 and RC-101 were prepared by solid-phase peptide synthesis, and the correct folding was ensured as described previously ^58-61^. The activity of HNP1 and RC-101 was confirmed using the recombinant ACD domain of *Vibrio cholerae* MARTX toxin ^42^. Recombinant SARS-CoV-2 Spike glycoprotein was purchased from R&D Systems (Minneapolis, MN). Bovine serum albumin (BSA) was purchased from Thermo Fisher Scientific (Waltham, MA). pCAGGS vector encoding SARS-CoV-2 Spike glycoprotein was obtained from BEI Resources (Manassas, VA; deposited by Dr. Florian Krammer, Icahn School of Medicine at Mount Sinai). Lentiviral NanoLuc expression vector (a gift from Erich Wanker; Addgene plasmid #113450; RRID:Addgene_113450) ^62^ was modified to introduce IL6 secretion signal (IL6ss) upstream of Myc-Nluc using PCR-based approach with NEBuilder DNA assembly (New England Biolabs, Ipswich, MA). Lentiviral helper plasmids expressing HIV Gag-Pol, HIV Tat, or HIV Rev under a CMV promoter were obtained from BEI Resources (Manassas, VA; deposited by Dr. Jesse Bloom, Fred Hutchinson Cancer Research Center).

### Cell lines

All cells were cultured at 37°C with 5% CO2 in a humidified incubator. U2OS and Vero E6 cells were grown in Dulbecco’s Modified Eagle Medium, H1299 in RPMI-1640 medium, Calu-3 in Eagle’s Minimum Essential Medium supplemented with 1% penicillin-streptomycin and 10% fetal bovine serum (FBS). H1299, Calu-3, Vero E6, and HEK293T cells were purchased from ATCC. The identity and purity of the U2OS cells were verified by STR profiling (Amelogenin + 9 loci) at the Genomic Shared Resource (OSU, Comprehensive Cancer Center) with 100% match using The Cellosaurus cell line database ^63^. All cell lines were mycoplasma-negative as determined by a PCR-based approach ^64^.

### Spike-mediated cell fusion assays

Using Lipofectamine 3000 reagent (Thermo Fisher Scientific, Waltham, MA), U2OS cells were co-transfected with pmCherry and pCAGGS vector encoding SARS-CoV-2 S protein. For a negative control, U2OS cells were transfected with pmCherry alone. 24-hours post-transfection, U2OS cells were trypsinized, mixed with Calu-3 cells on 96-well µ-plate (Ibidi, Germany), and incubated for 24 h in the presence and absence of HNP1 or RC-101 (40 µM; final concentration) with and without FBS (10%; final concentration). Following the incubation, the co-cultures were contra-stained with 1 µg/mL Hoechst 33342 (Invitrogen, Carlsbad, CA) and imaged using Nikon inverted microscope Eclipse Ti-E (Nikon Instruments, Melville, NY). Nikon NIS Elements AR software was used to quantify fusion indices calculated as number of nuclei per individual mCherry-positive syncytium.

### Fluorescence anisotropy

Increasing concentrations of SARS-CoV-2 Spike protein or BSA were incubated with 50 nM Cy5-labeled HNP1 in phosphate buffered saline (PBS). Fluorescence anisotropy was measured using a Tecan Infinite M1000Pro plate reader (Tecan SP, Inc., Baldwin Park, CA). *K*d values were determined by fitting the experimental data to the binding isotherm equation:

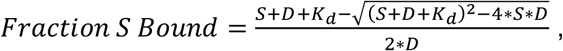

where *S* is the concentration of Spike (or BSA) and *D* is the concentration of defensin (Cy5-HNP1).

### Differential scanning fluorimetry (DSF)

DSF was performed as described previously ^42, 65^. Briefly, SARS-CoV-2 Spike protein was diluted to 1 µM in PBS in the absence or presence of HNP1 or RC-101 and supplemented with 1:5000 dilution of Sypro Orange dye (Invitrogen, Carlsbad, CA). Changes in fluorescence of the dye, which preferentially binds to protein hydrophobic regions exposed upon thermal-induced unfolding, were measured using a CFX Real-Time PCR Detection System (Bio-Rad, Hercules, CA). The melting temperatures (T_m_) were determined as the maximum of the first derivative of each normalized experimental curve.

### Pseudotyped virus production, cell infection, and luciferase assays

Virus-like particles pseudotyped with SARS-CoV-2 Spike protein were produced by transfecting HEK293T cells with pLenti-IL6ss-Myc-Nluc, pCAGGS-SARS-CoV-2 Spike plasmid, HIV Gag-Pol plasmid, HIV Tat plasmid, and HIV Rev plasmid using Fugene 6 transfection reagent (Promega, Madison, WI). Virus-containing medium was titrated to ensure undersaturating conditions for infection of H1299 cells. H1299 cells were incubated with the pseudovirus for 24 h followed by luciferase assays performed using NanoGlo Luciferase Assay System (Promega, Madison, WI). In one set of experiments, H1299 cells were pretreated with the defensins in the absence or presence of 10% serum before the addition of the pseudovirus; in the other set of exepriments, pseudovirus was preincubated with 50 µM defensins prior to addition to cells. Luminescence signals were recorded using a Tecan Infinite M1000Pro plate reader (Tecan SP, Inc., Baldwin Park, CA).

### SARS-CoV-2 infections and flow cytometry

SARS-CoV-2 strain USA-WA1/2020 provided by BEI Resources was plaque purified on Vero E6 cells to identify plaques lacking mutations in the Spike protein furin cleavage site. Non-mutated plaques were then propagated using Vero E6 cells stably expressing TMPRSS2 (kindly provided by Dr. Shan-Lu Liu, Ohio State University). Virus-containing supernatant aliquots were snap frozen in liquid nitrogen and stored at -80 °C. The virus stock was titered on Vero E6 cells. For infection experiments, virus at an MOI of 1 was used to infect Vero E6 in the presence or absence of FBS or defensins. After 24 h, cells were trypsinized, fixed with 4% paraformaldehyde for 1 h at room temperature, permeabilized with 0.1% Triton X100 in PBS, and stained with anti-SARS-CoV-2 N protein antibody (Sino Biological, Wayne, PA; #40143-MM08). Primary antibody labeling was followed by goat anti-mouse Alexa Fluor 647 secondary antibody (Life Technologies, Carlsbad, CA). Flow cytometry data acquisition was performed using a FACSCanto II flow cytometer (BD Biosciences, San Jose, CA). Data was analyzed using FlowJo software. Mock infected control samples were used to set gates for quantification of infection cell percentages.

Inhibitory concentration (IC_50_) of HNP1 was determined by fitting the data to the Hill equation:

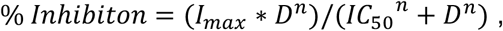

where *l*_*max*_ is the maximal inhibition, *D* is the concentration of defensin (HNP1), *n* is the Hill coefficient.

### Statistical analysis

Data are presented as mean values from several independent experiments (as indicated in the figure legends); error bars represent standard deviations (SD) or standard errors (SE) of the mean. Two-sample equal variance Student’s t-test with two-tailed distribution was used for data comparison; p-values <0.05 were considered statistically significant; individual p-values are indicated on the graph figures.

## AUTHORS CONTRIBUTIONS

EK: Conceptualization, Methodology, Validation, Formal analysis, Investigation, Writing - Original Draft, Writing - Review & Editing, Visualization

AZ: Investigation, Writing - Review & Editing GV: Investigation, Writing - Review & Editing

AS: Conceptualization, Validation, Writing - Review & Editing, Supervision, Funding acquisition WL: Resources, Writing - Review & Editing

JSY: Formal analysis, Resources, Writing - Review & Editing, Supervision, Project administration, Funding acquisition

DSK: Conceptualization, Methodology, Validation, Resources, Writing - Original Draft, Writing - Review & Editing, Supervision, Project administration, Funding acquisition

